# Cartilaginous and osteochondral tissue formation by human mesenchymal stem cells on three-dimensionally woven scaffolds

**DOI:** 10.1101/395202

**Authors:** Benjamin L. Larson, Sarah N. Yu, Hyoungshin Park, Bradley T. Estes, Franklin T. Moutos, Cameron J. Bloomquist, Patrick B. Wu, Jean F. Welter, Robert Langer, Farshid Guilak, Lisa E. Freed

## Abstract

The development of mechanically functional cartilage and bone tissue constructs of clinically relevant size, as well as their integration with native tissues, remain important challenges for regenerative medicine. The objective of this study was to assess adult human mesenchymal stem cells (MSC) in large, three dimensionally woven poly(ε-caprolactone) (PCL) scaffolds in proximity to viable bone, both in a nude rat subcutaneous pouch model and under simulated conditions *in vitro*. In Study I, various scaffold permutations: PCL alone, PCL-bone, “point-of-care” seeded MSC-PCL-bone, and chondrogenically pre-cultured Ch-MSC-PCL-bone constructs were implanted in a dorsal, ectopic pouch in a nude rat. After eight weeks, only cells in the Ch- MSC-PCL constructs exhibited both chondrogenic and osteogenic gene expression profiles. Notably, while both tissue profiles were present, constructs that had been chondrogenically pre-cultured prior to implantation showed a loss of glycosaminoglycan (GAG) as well as the presence of mineralization along with the formation of trabecula-like structures. In Study II of the study, the GAG loss and mineralization observed in Study I *in vivo* were recapitulated *in vitro* by the presence of either nearby bone or osteogenic culture medium additives but were prevented by a continued presence of chondrogenic medium additives. These data suggest conditions under which adult human stem cells in combination with polymer scaffolds synthesize functional and phenotypically distinct tissues based on the environmental conditions, and highlight the potential influence that paracrine factors from adjacent bone may have on MSC fate, once implanted *in vivo* for chondral or osteochondral repair.

## Introduction

Adult mesenchymal stem cells (MSCs) harvested from the bone marrow are potentially useful tools in the emerging field of regenerative medicine due to their relative ease of isolation, proliferation capacity [1], multipotency [2-4], and roles in modulating immune response [5] and neovascularization [6, 7]. MSCs have been used as a cell source for a variety of therapeutic approaches, including the repair of osteoarthritic lesions [8-12] and large bone defects caused by trauma or tumor resection [13, 14]. However, the development of mechanically functional cartilage and bone tissue constructs remains a continuing challenge.

Johnstone *et al*. [15] developed a protocol to induce chondrogenic differentiation of MSCs by *in vitro* exposure to insulin, selenous acid, ascorbic acid, pyruvate, dexamethasone, and transforming growth factor beta (TGF-β) that has been widely studied [3, 16-26]. In ongoing studies, a variety of environmental factors such as growth factors [27], mechanical stimulation [24, 28, 29], and hypoxia [30] have been shown to influence tissue engineered cartilage development from MSCs. While *in vitro* studies of MSCs have shown the potential for both chondrogenesis and osteogenesis, *in vivo* studies have shown that in some cases, chondrogenically-primed MSCs can develop a hypertrophic phenotype and exhibit mineralization [10, 17-20, 23, 31]. This tendency of MSCs toward hypertrophy has also been leveraged as a path toward bone tissue engineering [25], with defined contributions by both the donor MSCs and host cells to *in vivo* bone formation [26]. Similarly, other studies have shown production of biologically functional bone tissue containing marrow from MSCs and type I collagen scaffolds using a two-step process of chondrogenic priming followed by bone-priming with thyroxine and β-glycerol phosphate *in vitro* prior to implantation *in vivo* [32]. However, the environmental factors that lead to MSC hypertrophy in some cases and not in others are not fully understood.

The objective of the present study was to investigate various factors that influence chondral and mineralized tissue formation of MSCs seeded on three-dimensionally woven poly(*ε*-caprolactone) scaffolds (3D woven PCL) [19, 20]. We hypothesized that the pre- implantation culture conditions, as well as contact with a vital bone allograft to simulate *in vivo* engraftment, would influence MSC differentiation. The study was carried out in two phases: (Study I) an *in vivo* phase in which 3D woven PCL scaffolds (acellular, MSC-seeded, or chondrogenically pre-cultured MSC-seeded) were physically attached to viable bone grafts and implanted in an ectopic nude rat model [33] to evaluate the phenotype of tissue synthesized under these conditions; and (Study II) an *in vitro* study to observe the effect of soluble paracrine factors from viable bone on the differentiation of MSC-seeded scaffolds in a more controlled environment.

## Materials and Methods

### Cell Culture Media

*Control Culture Medium (CCM)* contained low glucose DMEM (Invitrogen, Thermo- Fisher Scientific, Waltham, MA) with 10% lot-selected fetal bovine serum (FBS; Atlanta Biologicals, Flowery Branch, GA), 10 ng/mL recombinant human fibroblast growth factor-2 (Peprotech, Rocky Hill, NJ) and 1% penicillin-streptomycin (P/S) (Invitrogen).

*Chondrogenic differentiation medium (Chondro)* consisted of high glucose DMEM (Invitrogen) containing 10 ng/mL recombinant human transforming growth factor beta 3 (rhTGF-β3; R&D Systems, Minneapolis, MN), 1% Insulin-Transferrin-Selenium (ITS+ Premix™; BD, San Jose, CA), 100 nM dexamethasone, 50 mg/L ascorbate-2 phosphate, 0.4 mM proline (Sigma-Aldrich, St Louis, MO) and 1% P/S (Invitrogen) [20].

*Osteogenic differentiation medium (Osteo)* consisted of low glucose DMEM containing 10% FBS, 1% P/S, 100 nM dexamethasone, 50 µM ascorbate-2 phosphate, and 10 mM beta glycerol phosphate (Sigma-Aldrich) [19].

### Human Mesenchymal Stem Cells

Undifferentiated MSCs were derived from bone marrow aspirates obtained from a healthy adult female donor at the Hematopoietic Stem Cell Core Facility at Case Western Reserve University. Informed consent was obtained, and an Institutional Review Board-approved aspiration procedure was used. Briefly, the bone marrow sample was washed with Dulbecco’s modified Eagle’s medium (DMEM) supplemented with 10% FBS. The sample was centrifuged at 460×*g* on a pre-formed Percoll density gradient to isolate the mononucleated cells. These cells were resuspended in serum-supplemented medium and seeded at a density of 1.8×10^5^ cells/cm^2^ in 10 cm diameter plates. Non-adherent cells were removed after four days by changing the medium. The adherent cells were expanded in CCM at 37°C in a humidified atmosphere of 5% CO_2_ [20]. The primary culture was trypsinized after approximately two weeks, cryopreserved using Freezing Medium (Invitrogen) for shipment and preparation of tissue engineered constructs.

### Tissue engineered constructs

Three dimensionally woven PCL scaffolds **(Fig. 1B)** were made as described previously [19, 20, 34], with some modifications to weaving parameters and scaffold architecture. Briefly, 156 µm diameter multifilament PCL yarns (EMS-Griltech, Domat, Switzerland) were woven in the x, y, and z directions to create a five-layered fabric measuring approximately 0.7 mm in thickness with an approximate porosity of 60%. The scaffolds were cut with an 8 mm diameter dermal biopsy punch and sterilized using ethylene oxide. Undifferentiated MSCs were expanded in CCM at 4,400 cells per cm^2^ through passage 2 (P2). The cells were trypsinized and then seeded on to the 3D woven scaffolds (1.25 million cells per scaffold) to produce tissue constructs.

**Figure 1.**
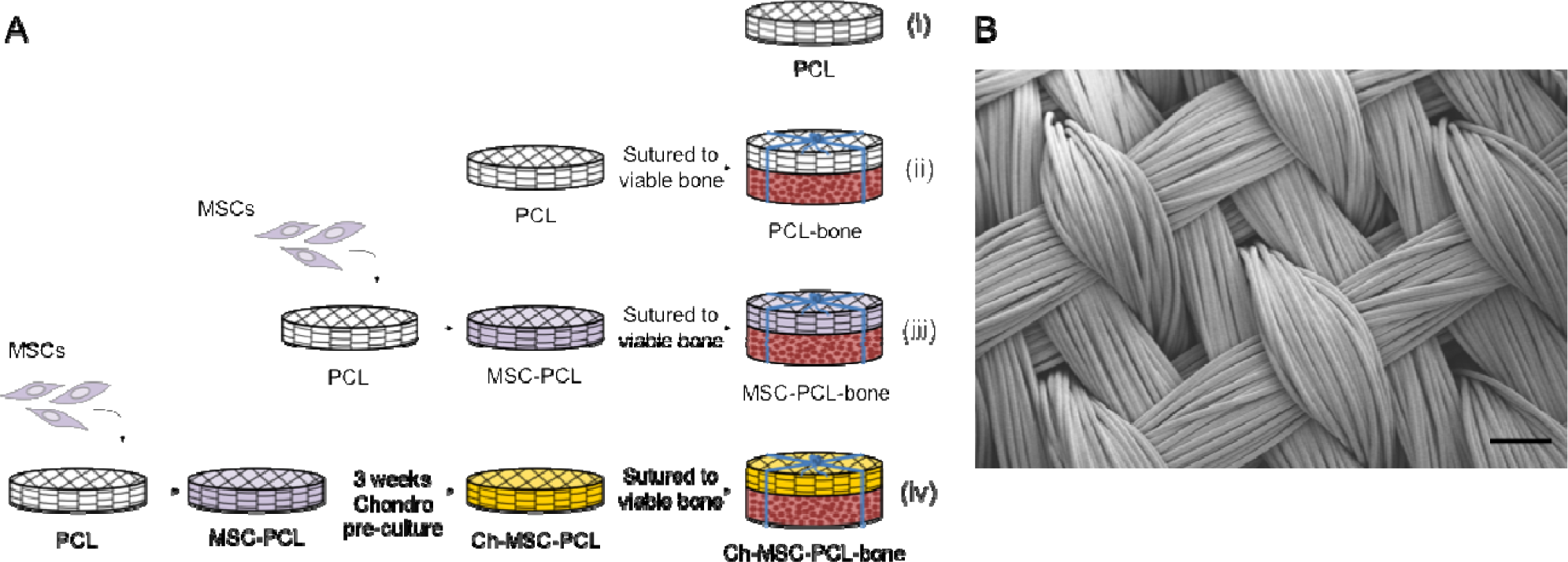
PCL scaffold and In Vivo Study Design Schema – Study I. (A) In vivo studies were carried out using (i) acellular PCL scaffolds (PCL), (ii) acellular PCL scaffolds sutured to bone (PCL-Bone), (iii) MSC- seeded scaffolds sutured to bone (MSC-PCL-Bone), (iv) and chondrogenically pre-cultured constructs sutured to bone (Ch-MSC-PCL-Bone) in a nude rat subcutaneous pouch model. (B) SEM image of 3D woven PCL scaffold. Scale bar = 200 µm.

### Bone and Cartilage Discs

The bone and cartilage discs were collected from femurs freshly harvested from two-week old bovine calves obtained from a local abattoir as described previously [33]. Briefly, 8 mm diameter osteochondral cores were harvested from the patellofemoral groove and sliced into 2 mm thick discs using a slotted cutting guide. Discs containing either trabecular bone only or cartilage only were used for the studies described herein; any disc containing a macroscopically visible combination of bone and cartilage was discarded. Bone and cartilage discs were gently washed in phosphate buffered saline and then cultured in CCM for up to three days prior to further testing.

### Study I: *In Vivo* Studies

An *in vivo* study was performed according to an Institutional Animal Care and Use Committee approved protocol that adhered to the NIH Guide for the Care and Use of Laboratory Animals and a previously established rodent model [33]. In brief, the constructs were implanted into two dorsal subcutaneous pouches on the backs of male immunodeficient (nude) rats weighing 180-240 grams (NIH RNU rats, Taconic Farms Inc, Germantown, NY) and were harvested after eight weeks.

The four specimen groups were: (i) acellular PCL scaffold (PCL), (ii) acellular PCL scaffold sutured to bone (PCL-bone), (iii) MSC-seeded PCL scaffold sutured to bone (MSC- PCL-bone), and (iv) chondrogenically pre-cultured MSC-seeded PCL scaffold sutured to bone (Ch-MSC-PCL-bone) (**Fig. 1A**). For MSC-seeded scaffolds (group iii), MSCs (P2) were seeded into PCL scaffolds and implanted after one to two days of culture in CCM. For Ch-MSC-PCL constructs (group iv), MSCs (P2) were seeded onto PCL scaffolds and subsequently cultured in Chondro medium (one construct and 5 mL of medium per 35 mm diameter well of a 6-well plate) for 3 weeks and then implanted. In Groups ii, iii, and iv, construct/bone composites were prepared following any requisite culture by suturing scaffolds to bone discs using 5.0 Vicryl (polyglycolide-co-lactide 90/10, Ethicon, Somerville, NJ). The suture was wrapped around the two discs and square knotted, wrapped around the two discs again perpendicularly to the first wrap, and knotted again. Two additional specimen control groups were added: bovine calf cartilage discs sutured to bone, and two bovine calf cartilage discs sutured together, following the same methods.

All specimens were maintained at 37°C in a standard incubator until four hours prior to implantation, at which time they were transferred to the operating room in a 4°C solution of Hank’s balanced saline (Invitrogen). Two constructs were implanted per rat, oriented with the bone positioned adjacent to the dorsal muscle and PCL-based construct positioned adjacent to the skin. A total of 26 rats were studied for a total of (n=6 rats/group + the two additional control rats). All rats were sacrificed after 8 weeks, for analysis of the constructs.

### Study II: *In Vitro* Model System

Additional studies were performed in an *in vitro* model system to observe the effect of soluble paracrine bone factors emanating from bone in proximity to the constructs that had been freshly seeded with MSCs (**Fig. 2A**) and those that had been chondrogenically pre-cultured (**Fig. 2B**). Constructs were cultured under various conditions in 6-well transwell inserts with 0.4 µm track-etched PET membranes (EMD Millipore). To increase soluble factor transport, four, 2 mm diameter holes were made in the walls of the inserts using a soldering iron. The inserts were then washed with deionized water and re-sterilized with ethylene oxide prior to use. Bone explants (*i.e.*, four quarters of an 8 mm diameter, 2 mm thick bone disc) were either absent or present in the well of the 6-well plate, thus the designation “nearby.”

**Figure 2.**
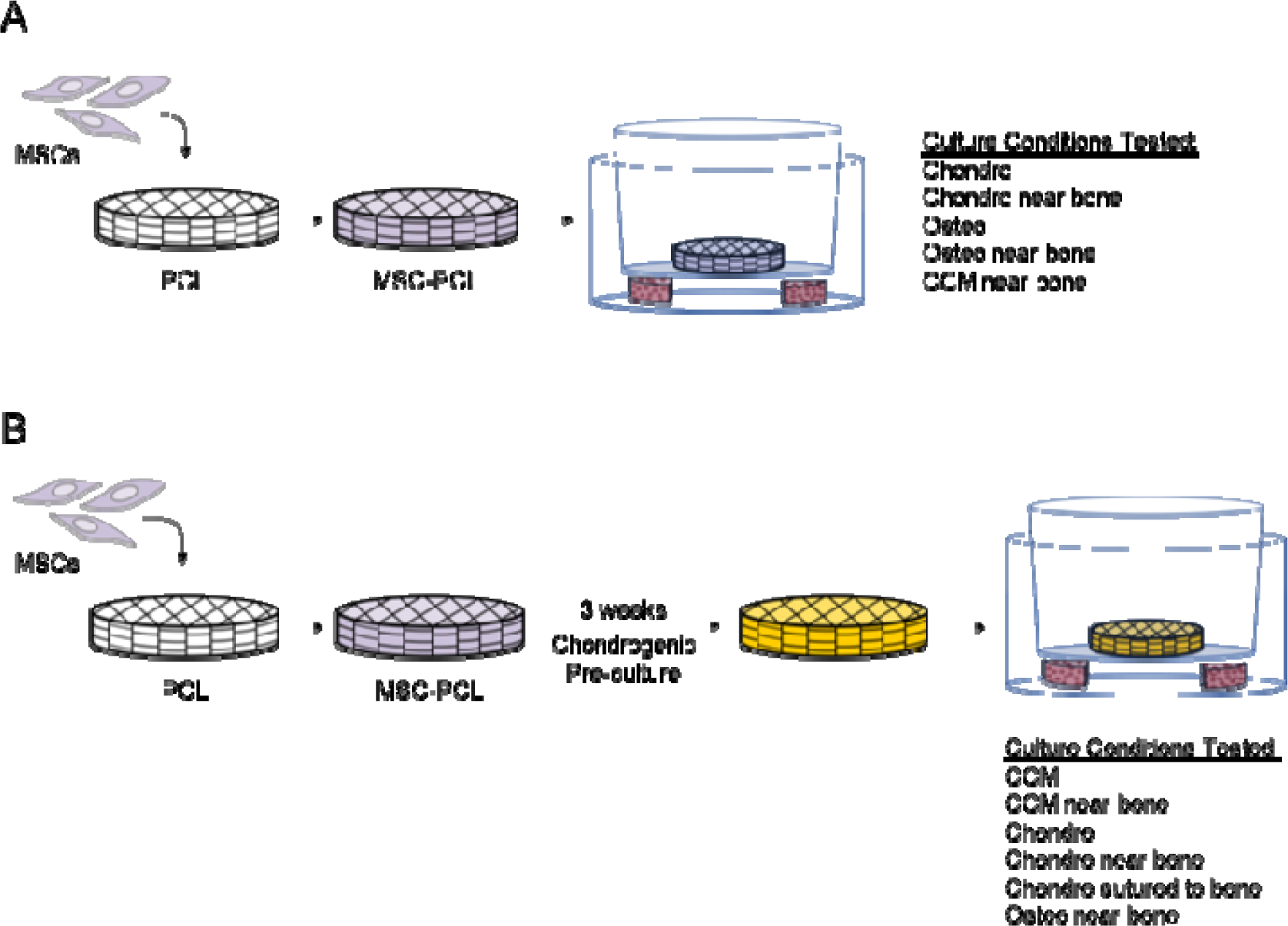
PCL scaffold In Vitro Study Design Schema – Study II. (A) Transwell plates were used to culture freshly seeded implants (MSC-PCL) in vitro in proximity of viable bone using various culture media. (B) In vitro culture was also carried out to recapitulate certain conditions of chondrogenically pre-cultured constructs in vitro with viable bone and various culture conditions.

Freshly seeded constructs (**Fig. 2A**) were studied under five culture conditions: Chondro medium without nearby bone, Osteo medium without nearby bone, Chondro medium with nearby bone, Osteo medium with nearby bone, and CCM with nearby bone. Constructs (n=4 per condition) were cultured for 3 weeks.

To recapitulate the conditions of chondrogenically pre-cultured implants *in vitro*, MSC-seeded constructs were first cultured in Chondro medium for three weeks as with the *in vivo* samples (**Fig. 2B**). Following the chondrogenic pre-culture, Ch-MSC-PCL samples were placed into one of six conditions (n=4 constructs/condition): Chondro medium without nearby bone, CCM without nearby bone, CCM with nearby bone, Osteo medium with nearby bone, Chondro medium with nearby bone, and Chondro medium in direct contact with bone. In the direct contact group, the pre-cultured constructs were sutured directly to bone discs as in the *in vivo* study and cultured in standard 6-well plates.

### Mechanical Testing

In Study I, all implants were harvested after 8 weeks *in vivo*, wrapped in saline soaked gauze and frozen at -20°C until testing. Specimens were thawed at the time of testing, and the aggregate moduli of explants were quantified by radially confined compression testing as described previously [19, 20, 34]. In brief, after removing the lower bone component, when present, 3 mm diameter specimens were cored from each explanted construct and tested using an ELF 3200 Series precision controlled materials testing system (Bose, Framingham, MA).

### Cell, Molecular, and Biochemical Analyses

Following mechanical testing, the specimens were quartered and the sample was subjected to molecular analyses as follows. RNA was isolated from the samples by RNeasy kit (Qiagen, Germantown, MD), and quantitative (qPCR) was performed with an RNA-to-Ct™ kit using gene specific, validated TaqMan primers listed as follows (Applied Biosystems, Foster City, CA): *ACAN* (aggrecan): Hs00153936_m1, *BSP* (bone sialoprotein): Hs00173720_m1, *COMP* (cartilage oligomeric matrix protein): Hs00164359_m1, *COL2A1* (collagen type 2a1): Hs00264051_m1, *OC* (osteocalcin): Hs00958189_m1, *RUNX2* (runt-related transcription factor 2): Hs00231692_m1, *SP7* (osterix): Hs01866874_s1, *SOX9* (SRY (sex determining region Y)- box 9): Hs01001343_g1. The chondrogenic markers assayed were *SOX9, ACAN, COMP*, and *COL2A1*, the osteogenic markers assayed were *BSP*, *RUNX2*, *OC*, and *SP7*. All values were normalized to human *GAPDH* and again to P2 undifferentiated MSCs.

Sequencing Flowcell cluster amplification for RNA-seq was performed according to the manufacturer’s recommendations using the V3 TruSeq PE Cluster Kit and V3 TruSeq Flowcells (Illumina, San Diego, CA). Flowcells were sequenced with 101 base paired end reads on an Illumina HiSeq2000 instrument, using V3 TruSeq Sequencing by synthesis kits and analyzed with the Illumina RTA v1.12 pipeline (Illumina). An index of human stem cell characteristics, Rohart’s score [35, 36], was determined by analyzing transcriptome profiles with Stemformatics tools. Briefly, data were uploaded to Stemformatics.org, an academic bioinformatics tool for data visualization and analysis. Rohart’s test was performed on transcriptome data to determine likelihood of MSC-like characteristics in unknown samples versus hundreds of known MSC and non-MSC transcriptomes existing in the database. Samples with a Rohart score above 0.6 are considered MSCs. The raw data used to generate the Rohart scores were accessed from the National Center for Biotechnology Information Gene Expression Omnibus database (GEO accession number will be available after acceptance of manuscript).

RNAseq reads were preprocessed to remove Nugen adapters using Trimomatic version 0.32. Trimmed reads were then aligned with bowtie version 1.01 to a target containing the bovine (bt8), rat (rn4) and human (hg38) genomes with a custom gene annotation file consisting of the longest transcript at each locus for each genome and gene expression was then quantified using RSEM version 1.2.15. The contribution from each species is expressed as the sum of RSEM counts assigned to each species divided by the total counts. (The raw sequencing files, combined genome target, custom annotation file and counts per gene will be deposited in the Gene Expression Omnibus upon acceptance of the manuscript).

To quantify DNA and GAG, approximately 1/4 of each sample was minced followed by digestion with papain. DNA content was measured by fluorescence using the Picogreen assay kit, and sulfated GAG content was measured using the Blyscan assay kit (Biocolor Life Sciences, Carrickfergus, UK). Calcium content was measured using a colorimetric kit (Biovision, Milpitas, CA) as previously described [19]. The GAG and calcium data were subsequently normalized by specimen wet weight.

### Histology analysis

The remaining portions of each sample were fixed in 10% neutral buffered formalin (Sigma-Aldrich) and placed at 4°C for at least 24 hours. Samples were then embedded in paraffin and partially decalcified (Immunocal solution, Decal Chemical Corporation, Suffern, NY) for 10 min at room temperature (RT) for sectioning. Ten micron thick sections were placed on slides and stained with hematoxylin and eosin (H&E), safranin O/fast green, or von Kossa/nuclear fast red. For immunostaining, paraffin section slides were de-paraffinized in reveal solution in a pressure cooker for 20 min. Slides were then incubated with 10% horse serum for 1 hour at RT, primary mouse anti-human antibody against HSNA (clone 235-1, EMD Millipore) at 1:100 dilution overnight at 4°C, followed by secondary goat-anti-mouse antibody Alexa488 (Invitrogen) at 1:100 dilution for 30 min at RT. Control slides were processed identically but without primary antibody. All slides were then mounted with Vectashield containing DAPI (Vector Laboratories, Burlingame, CA) and coverslipped.

### Microcomputed Tomography (microCT)

MicroCT imaging was performed as described previously [19]. Briefly, specimens were formalin fixed, immersed in 70% ethanol, and imaged using a microCT scanner (MicroCT 40, Scanco Medical, Brüttisellen, Switzerland). Medium-resolution scans were obtained in 16 µm increments at energy of 45 kVp and intensity of 177 μA. Three-dimensional reconstructions were obtained using accompanying commercial software (Scanco).

### Statistical Analysis

Data were calculated as mean ± standard error and analysed using multi-way analysis of variance (ANOVA) in conjunction with Tukey’s HSD or Dunnet’s *post hoc* test (*α* =0.05) using Statistica (v.7, Tulsa, OK). A sample size of n ≥ 3 was used for all assays. Values of *p <0.05* were considered statistically significant.

## Results

### Study I

Ch-MSC-PCL constructs, which had been chondrogenically pre-cultured for 3 weeks, contained a high level of GAG (10.76 ± 0.46 µg/mg wet wt.) and no detectable calcium (**Fig. 3**) prior to implantation. Histological cross sections of these implants also showed positive staining for GAG and negative staining for mineral (**Fig. 4**). After 8 weeks implantation *in vivo*, tissue incorporation and development were observed in all constructs, regardless of pre-culture or MSC seeding, as evidenced by the presence of dsDNA at 6-8 ng/mg construct (**Fig. 3A**) and histological analysis demonstrating complete filling of the pore structure of the scaffolds with the presence of counterstained nuclei (**Fig. 4**). All scaffolds that were sutured to a viable bone graft showed excellent integration between the scaffold and the bone, with neotissue bridging between the 3D woven PCL and the native bone trabeculae (**Fig. 4**).

**Figure 3.**
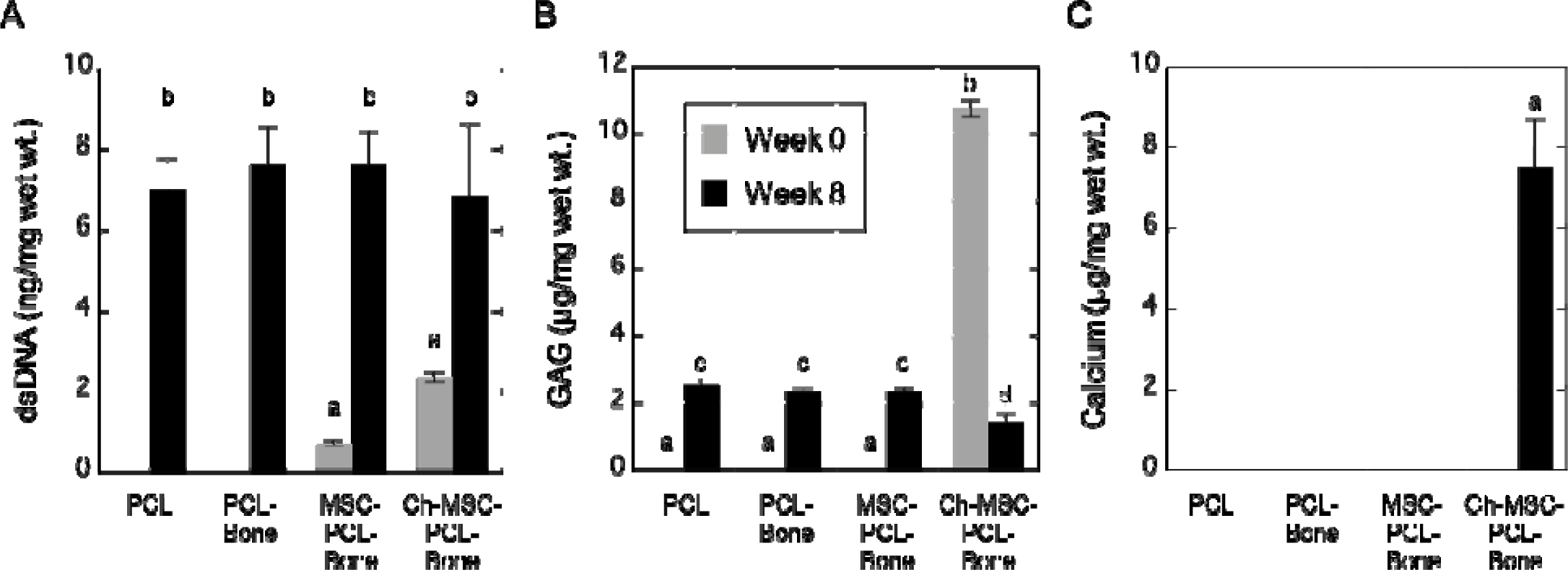
Biochemical characterization of in vivo implants. (A) Total DNA (ng/mg wet weight); (B) GAG content (µg/mg wet weight); and (C) calcium content (µg/mg wet weight). Data from time-zero implants and those harvested after 8-weeks in vivo appear gray and black, respectively. Groups not sharing the same letters are statistically different (p < 0.05).

**Figure 4.**
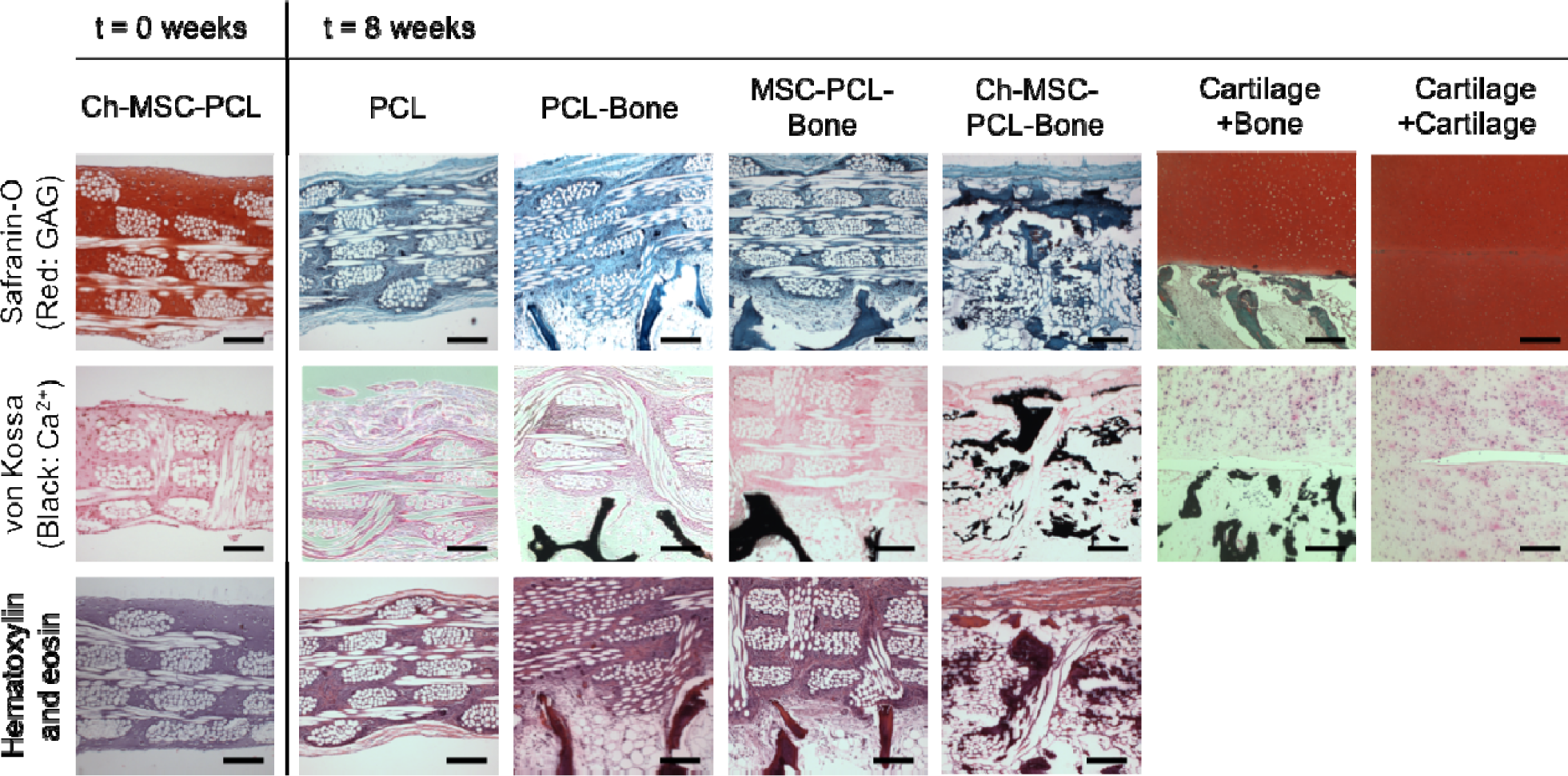
Histological results. Representative histological sections for Ch-MSC-PCL prior to implantation (positive control) and all implants following 8 weeks in vivo with additional control tissue sections for bovine cartilage and bone controls following 8 weeks in vivo. Tissues were treated with safranin-O/fast green (orange-red color indicates presence of GAG; the PCL fibers and fiber bundles appear white within the constructs), von Kossa staining (black color indicates presence of mineralized matrix), and H&E staining. Scale bars represent 200 µm.

No statistical difference was noted in dsDNA content across all groups (p>0.05). With respect to structure and extracellular matrix (ECM) composition, a GAG content of approximately 2 μg/mg wet weight was detected by biochemical assay in groups not undergoing pre-culture (PCL, PCL-bone, and MSC-PCL-bone) **(Fig. 3B)**. However, no safranin O staining was evident in these specimens **(Fig. 4)**. No calcium was detected in these groups by biochemical analysis (**Fig. 3C**) or histological staining **(Fig. 4)**.

In contrast, the Ch-MSC-PCL constructs, after 8 weeks *in vivo*, contained approximately 7 μg/mg of calcium **(Fig. 3C**), and had lost more than 80% of their initial GAG (**Fig. 3B**), with negative Saf-O/Fast Green histological staining for GAG and positive von Kossa histological staining for mineral **(Fig. 4)**. MicroCT imaging of the Ch-MSC-PCL constructs showed mineral deposition in a trabecular pattern, while no mineral was observed in groups that were not pre-cultured in Chondro medium **(Fig. 5A)**. No mineralization was present by histological staining in an additional control group where bovine calf cartilage was sutured to vital bone, implanted in nude rats and harvested after eight weeks **(Fig. 4).** The aggregate modulus of constructs following 8 weeks *in vivo* was significantly higher for the Ch-MSC-PCL-bone group (1.31 ± 0.56 MPa) as compared with either the MSC-PCL-bone (0.319 ± 0.046 MPa), PCL-bone (0.36 ± 0.40 MPa) or PCL (0.26 ± 0.018 MPa) group (**Fig. 5B**).

**Figure 5.**
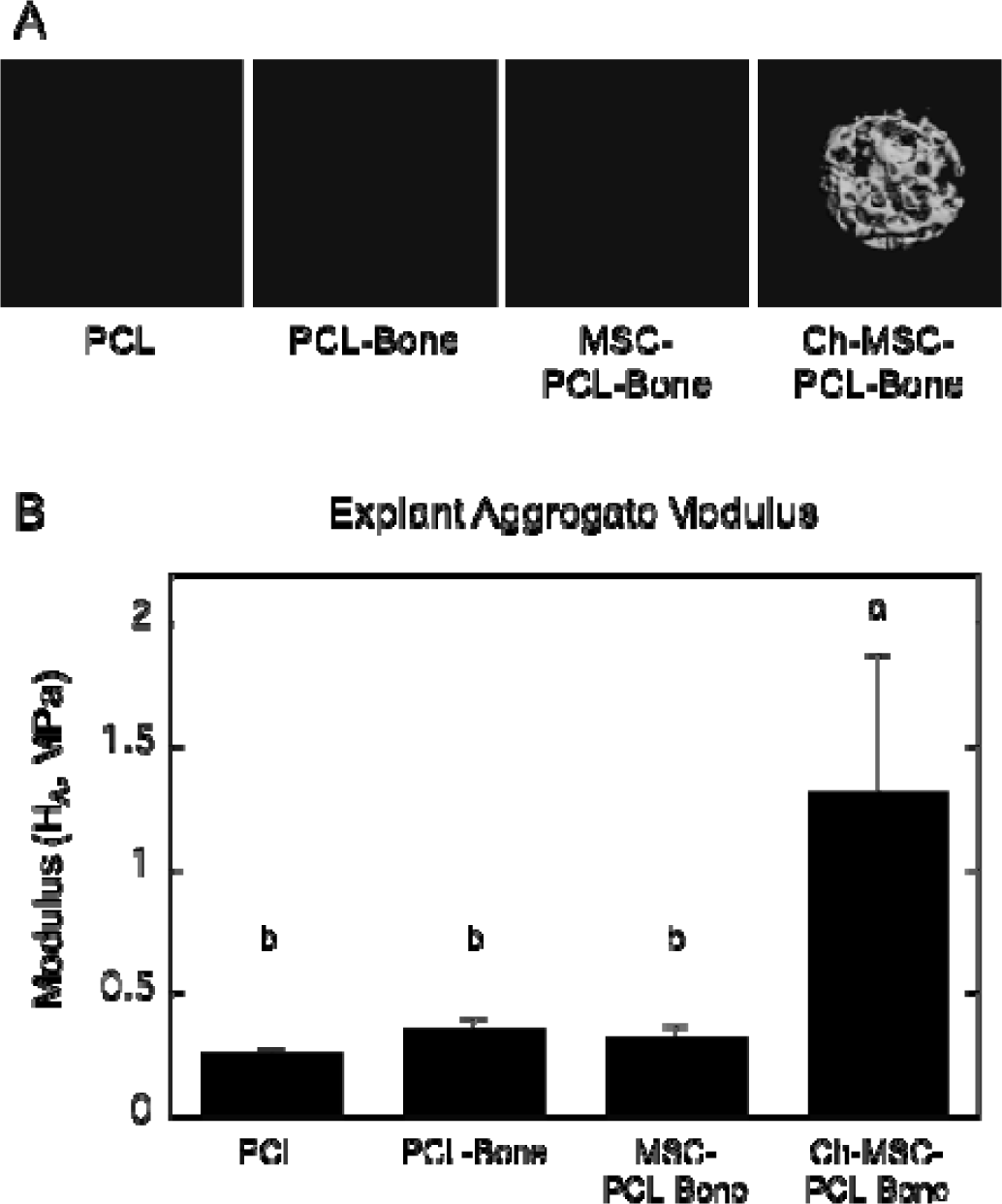
Microcomputed tomography and mechanical properties. (A) microCT images and (B) Aggregate moduli (H_A_) of the upper components of harvested PCL, PCL-bone, MSC-PCL-bone, and Ch-MSC-PCL- bone constructs. Groups not sharing the same letters are statistically different, (p < 0.05).

The Rohart score is a method to classify mesenchymal stem cell properties based on transcriptome data, with values above 0.60 considered stem cells. Prior to any pre-culture, MSCs had a high Rohart score of ∼0.85. Ch-MSCs had a low Rohart score of -0.26 following chondrogenic pre-culture prior to implantation, and a score of 0.11 after 8 weeks *in vivo*.

At the time of implantation, the Ch-MSC-PCL group expressed chondrogenic marker genes (*SOX9, ACAN, COMP*, and *COL2A1*) at high levels, whereas osteogenic markers (*BSP, RUNX2, OC* and *SP7*) were absent or nearly absent (**Fig. 6**). In contrast, at the time the constructs were harvested, the Ch-MSC-PCL group maintained high levels of chondrogenic markers but also expressed significantly higher levels of osteogenic markers (**Fig. 6**). For the chondrogenic markers, *ACAN* demonstrated the highest relative increase from week 0 to week 8 with an increase in gene expression of 3 orders of magnitude. *COL2A1* also increased significantly but only changed one order of magnitude relative to the time zero value. For the osteogenic markers, *BSP* increased approximately 2 orders of magnitude; whereas *OC* remained unchanged after 8 weeks *in vivo* (**Fig. 6**).

**Figure 6.**
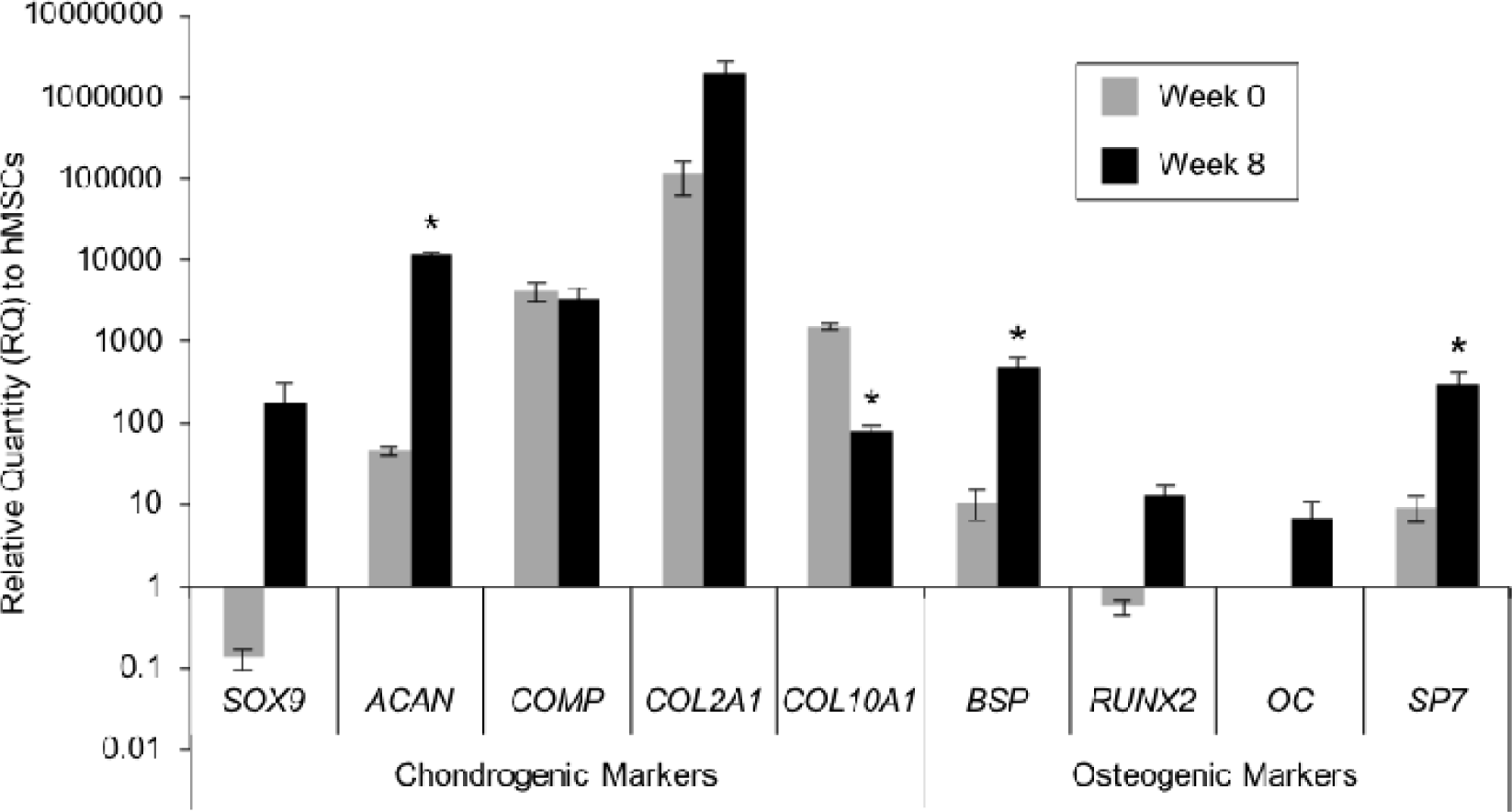
Gene expression in the Ch-MSC-PCL group. Human gene expression, shown as a relative quantity (RQ) to MSCs, for chondrogenic and osteogenic markers. Data from time-zero implants and those harvested after 8-weeks in vivo appear gray and black, respectively. * indicates p < 0.05 as compared to gene expression at 0 weeks.

Following harvest, human DNA could only be visualized in the constructs from the Ch- MSC-PCL-bone group using human specific nuclear antigen (HSNA) (**Fig. S1 A**). In the harvested Ch-MSC-PCL-bone group, transcriptome analyses revealed that human RNA represented ∼45% of the total RNA in the Ch-MSC group, with the balance consisting mainly of rat RNA with traces of bovine calf RNA (**Fig. S1 B**).

### Study II

To investigate potential mechanisms responsible for GAG loss and mineralization observed in the Ch-MSC explants, some aspects of the *in vivo* environment were recapitulated *in vitro* in order to study specific roles of the culture medium formulation and soluble paracrine factors released from the adjacent bone grafts. Studies were carried out to observe the effects of these variables on freshly seeded constructs **(Fig. 2A)** and, similar to Study I, on chondrogenically pre-cultured constructs (Ch-MSC-PCL) (**Fig. 2B**).

In freshly seeded constructs, histological analysis (Saf-O/FastGreen) indicated GAG accumulation in constructs cultured in Chondro media (with or without adjacent bone), but GAG was not detectable in constructs cultured in Osteo or CCM (**Fig. 7)**. Further, compositional analysis indicated that for constructs cultured in Chondro medium, the presence of vital bone in the transwell culture led to a significantly lower GAG content compared to the constructs cultured without the presence of bone (p=0.048) **(Fig. 7)**. Both histological and compositional analyses indicated calcium deposition when MSC-PCL constructs were cultured in osteogenic medium, with the presence of nearby vital bone significantly increasing the calcium content almost 2-fold over Osteo media alone (p<0.002) (**Fig. 7)**.

**Figure 7.**
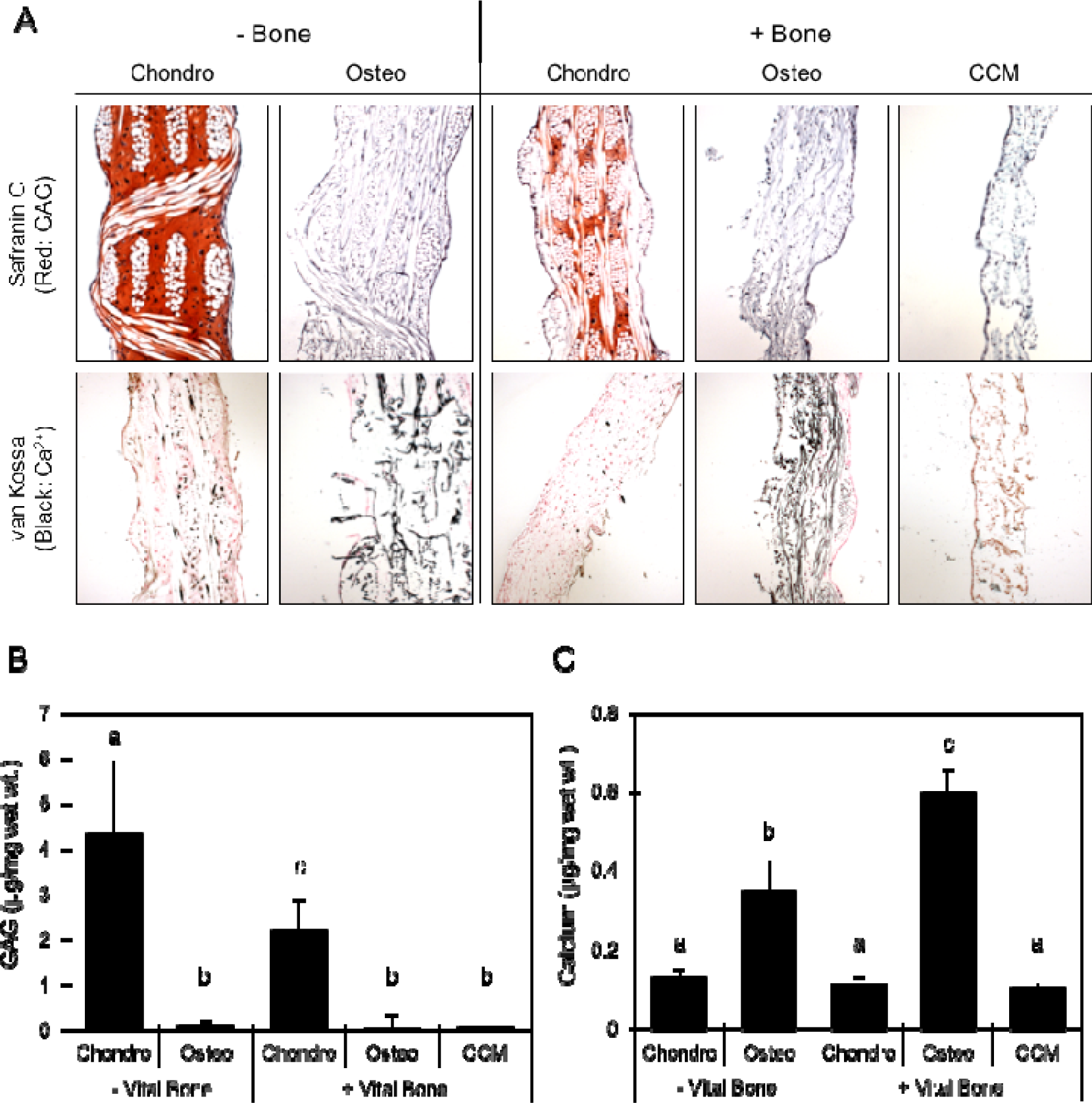
Freshly seeded MSC constructs cultured in vitro under various conditions. (A) GAG and mineral are shown histologically after staining with safranin O/fast green and von Kossa following 3 weeks in various culture conditions. (B) GAG and (C) calcium content of constructs following 3 weeks in various conditions are shown graphically (µg/mg wet wt.). Error bars represent one standard deviation. Groups not sharing the same letters are statistically different (p < 0.05)

GAG content increased significantly in the Ch-MSC-PCL constructs over the time between the first 3 weeks of chondrogenic pre-culture and the additional 3 weeks the constructs remained in the Chondro medium without the presence of bone **(Fig. 8 B)**. In contrast, when constructs that were chondrogenically primed were switched to CCM and cultured for an additional 3 weeks, GAG content decreased to 44% of its value at the end of chondrogenic preculture. GAG content plateaued in groups that were cultured in Chondro medium with either bone directly sutured to the construct or with bone in the wells of the transwell plate. In addition, GAG content decreased by approximately 50% when the culture condition of chondrogenically primed constructs was changed to either CCM or Osteo medium with nearby bone **(Fig. 8)**.

**Figure 8.**
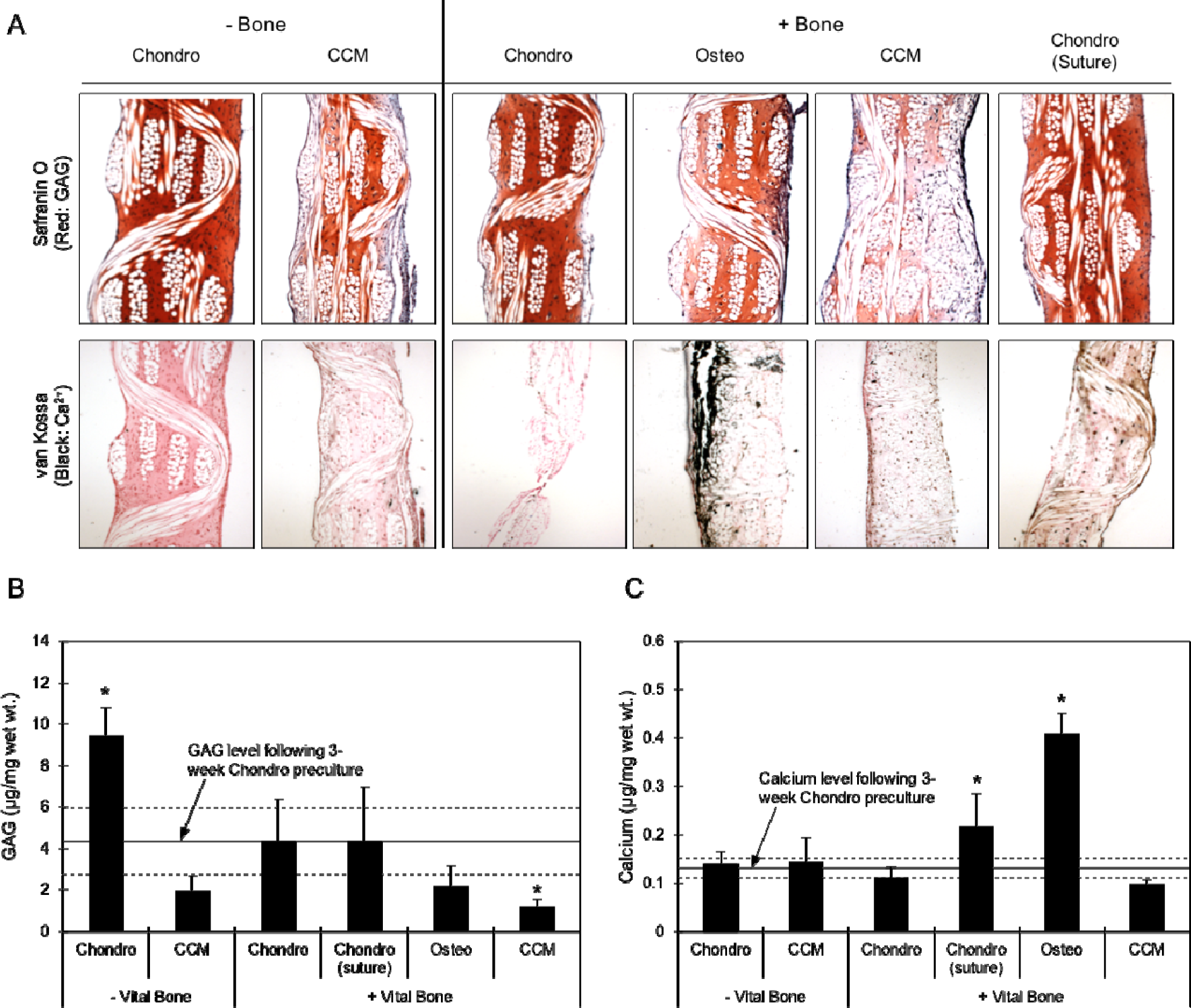
Ch-MSC-PCL constructs that were chondrogenically pre-cultured for three weeks were cultured for an additional three weeks in vitro under various conditions. (A) GAG and mineral are shown histologically after staining with safranin O/fast green and von Kossa. Measured contents of (B) GAG and (C) calcium. Shaded area represents the average value following initial 3-week chondrogenic pre-culture ± one standard deviation; * indicates p < 0.05 compared to value following initial 3-week chondrogenic pre-culture by one-way ANOVA with a Dunnet’s post-hoc test

Following the three-week chondrogenic pre-culture, calcium content was significantly increased when the constructs were cultured in Osteo medium with nearby bone and Chondro medium with sutured bone to values of ∼0.4 μg/mg wet wt. and ∼0.2 μg wet wt. respectively **(Fig. 8C)**. Regardless of the presence of bone in the transwell plate, incubation in CCM and Chondro media resulted in no significant changes in the mineralization of the constructs compared to the end of the three-week pre-culture. Histological analysis indicated calcium deposition and surface mineralization when Ch-MSC-PCL constructs were incubated in the presence of bone, but not in the continued presence of chondrogenic medium additives and absence of bone (**Fig. 8A**).

## Discussion

The findings of this study show the capacity for cartilaginous or mineralized tissue synthesis by MSCs in 3D woven PCL scaffolds, both *in vivo* and *in vitro* [20, 34], and provide important information on the effects that viable bone and culture conditions have on the structure, composition, and mechanical properties of the resulting engineered tissue. First, the *in vivo* tissue development and mechanical characteristics assessed 8 weeks following ectopic implantation in the back of a nude rat showed that pre-culture conditions followed by contact with viable bone *in vivo* could significantly shift the phenotype of MSCs from a chondrogenic phenotype to a hypertrophic, osteogenic one. Next, to further ascertain the influence soluble factors in proximity to viable bone on differentiation, Study II was carried out *in vitro* to simulate aspects of the *in vivo* environment, and confirmed the importance of culture medium composition and the presence of nearby or adjacent bone in modulating MSC chondrogenesis and osteogenesis. These findings provide new insights into the mechanisms by which MSCs – in some cases – undergo a shift to a hypertrophic phenotype once implanted for chondral or osteochondral repair [10, 17-20, 23, 31].

The results of our study indicate that PCL scaffolds support cellularization as well as extracellular matrix synthesis, accumulation, and remodeling *in vivo*, regardless of whether or not MSCs were pre-seeded or pre-cultured. These constructs yielded a repair tissue with hybrid properties that in some cases resembled cartilage and/or bone, depending on the environment (*in vitro* or *in vivo*), culture conditions (Chondro/Osteo/CCM), and the presence of bone in proximity to the construct. In our *in vivo* model, this finding was noted in all groups, suggesting that the 3D woven PCL scaffold was highly conducive to cell infiltration and *de novo* tissue synthesis. In terms of clinical translation of 3D woven scaffolds for osteochondral tissue repair, the data suggest that an acellular PCL implant may support rapid endogenous cell infiltration without the need for autologous cell harvest or *ex vivo* cultured cells. After 8 weeks in an ectopic model, constructs from the PCL, PCL-bone, and MSC-PCL-bone groups had no detectable calcium and low levels of GAG (2 μg/mg wet wt.), but aggregate moduli that were within the reported values for normal articular cartilage (H_A_ of 0.1-2.0 MPa) [37, 38]. This observation is consistent with our previous *in vitro* studies, which reported the generation of mechanically functional cartilage *in vitro* through mechanical and physical interactions with the biomimetic scaffold [39, 40].

In contrast, following 8 weeks in a rat model, constructs from the Ch-MSC-PCL-bone group had calcium deposits, very small amounts of GAG, and aggregate moduli of 1.5 MPa, which is approximately five-fold higher than the aforementioned three groups. The Ch-MSC- PCL-bone constructs had clearly undergone substantial remodeling *in vivo* since at the time of implantation, they were characterized by significant markers of chondrogenesis (e.g., abundant type II collagen and GAG. Consistent with this observation, the Ch-MSC-PCL-bone constructs expressed both chondrogenic and osteogenic genes both after the *in vitro* pre-culture period and at the time of harvest. The observed *in vivo* shift of chondrogenically primed constructs towards a hybrid osteochondral phenotype with characteristics of both cartilage and bone adds to the growing body of evidence that chondrogenically primed MSCs can assume a hypertrophic phenotype with upregulated osteogenic gene expression [19, 26, 32].

With respect to this hypertrophy, it is well established that the use of TGF-β3 for *in vitro* chondrogenesis of MSCs results in the presence of hypertrophic markers and may lead to bone formation [22, 41-44]. Indeed, markers such as type X collagen, MMP13, and alkaline phosphatase indicate a hypertrophy consistent with endochondral ossification observed during skeletal development [43]. The hypertrophy and ECM mineralization that was observed in the current study are consistent with that observed by others in similar ectopic studies [22, 41, 42, 45]. Using a pellet culture system, a previous study has demonstrated significant accumulation of calcified matrix after MSC derived pellets had been chondrogenically induced using TGF-β3 prior to ectopic implantation in a SCID mouse, which was not observed with human articular chondrocyte derived pellets [22] and also not observed with our bone-cartilage controls. Additionally, as with the current study, calcification of MSCs was not observed when cultures were maintained *in vitro* under the same chondrogenic conditions.

Along these lines, recent studies have provided insights into the signaling pathways and expansion conditions that potentially predispose MSCs to hypertrophy and endochondral ossification. For example, passaging the cells in bFGF containing expansion media pushed the cells towards endochondral ossification, whereas culturing with WNT3A in addition to bFGF suppressed the propensity for hypertrophic maturation [45]. The authors from this study concluded that the use of WNT3A and bFGF maintain the cells’ osteogenic and chondrogenic potential in subsequent differentiation protocols [45]. As it has been established that Wnt and hedgehog signaling are involved in regulating the differentiation of osteoprogenitor cells, the involvement of these signaling pathways and their roles in the endochondral ossification “switch” for MSCs can be anticipated [46-49]. More recently, using developmental paradigms, researchers have sought to elucidate the mechanisms of the Wnt/β-catenin pathway to regulate the fate of MSC progenitors undergoing osteoblastic and chondrogenic differentiation [50-52]. Inhibition of Wnt signaling during *in vitro* chondrogenesis significantly enhanced the synthesis of cartilage specific macromolecules in early culture [52, 53]. Similarly, a reduction in β-catenin was also shown to correlate with lower levels of the hypertrophic marker, type X collagen [54]. The emerging data on MSC differentiation suggest a complex signaling cascade wherein high levels of Wnt signaling inhibit chondrogenic differentiation while promoting hypertrophy [50, 51]. However, low levels of Wnt signaling are required to promote MSC chondrogenesis and inhibit the endochondral ossification phenotype that ultimately leads to calcification [50, 51]. These data indicate a more precise *in vitro* differentiation protocol, involving Wnt signaling, to stabilize and protect the chondrogenic profile of these cells and *de novo* tissues once implanted.

Other studies have also pointed to the potential use of the endochondral ossification pathway in generating bone constructs following chondrogenic induction. More specifically, the generation of intramembranous ossification of MSCs *in vivo* has remained challenging, and researchers have turned to the field of “developmental engineering” for bone formation [25, 55-57]. The prevailing hypothesis in these studies is that promoting endochondral ossification via chondrogenic hypertrophy will stimulate vascular endothelial growth factor (VEGF), which is important to support blood vessel formation to nourish the bone tissues and to ultimately avert the possibility of avascular necrosis and core degradation often observed in bone tissue engineering studies [56, 57]. Consistent with the current study, Scotti et al. demonstrated concomitant increases in type X collagen and BSP initiates endochondral ossification by MSCs and that implantation of the resulting constructs ectopically resulted in bone formation via an endochondral ossification pathway [55].

In the present study, the use of a transwell *in vitro* culture system of MSC-based constructs enabled the study of soluble biological cues without the influences of mechanical forces, host systemic responses, or animal-to-animal variability that can result in difficulties in interpreting *in vivo* studies. Specifically, introduction of specific Osteo medium additives along with viable bone to the chondrogenically primed constructs induced an osteogenic phenotype. The data showed that a chondrogenic phenotype can be maintained although the continued presence of chondrogenic medium was required, consistent with previous reports [12, 23, 58]. However, as previously discussed, the presence of hypertrophic genes following chondrogenic induction, namely *COL10A1* and *BSP*, suggest a shift to a hypertrophic phenotype upon removal of chondrogenic medium and *in vivo* implantation. It is also important to note that the use of serum, which was added to the Osteo and CCM media, may have contributed to the observed GAG loss in those groups due to the presence in serum of GAG-degrading enzymes. Nonetheless, the resulting phenotype for freshly seeded constructs *in vitro* was largely determined by the culture medium. Constructs cultured in Chondro medium had higher GAG content compared to those cultured in CCM and Osteo media.

The presence of viable bone had significant effects on the constructs, and in the Chondro culture, resulted in constructs with lower GAG content compared to Chondro medium alone. Conversely, when cultured in Osteo medium, constructs had higher calcium content compared to other culture conditions. The addition of the viable bone to the Osteo culture system increased the calcium deposition by an additional 71% compared to Osteo media alone, suggesting paracrine factors influencing the fate of the differentiating cells. In short, the GAG loss and ECM mineralization that were observed *in vivo* in the Ch-MSC-PCL-bone group were recapitulated in transwell plates, by maintaining viable bone in close proximity to the Ch-MSC-PCL constructs and allowing soluble bone-derived factors to influence the cells. This upregulation of osteogenic markers due to soluble factors from bone is consistent with a previous study of chondrocyte- agarose constructs [59]. Furthermore, the data are consistent with other reports on cell co-culture and associated paracrine signaling that have been shown to drive MSC differentiation [29, 58, 60-64]. While we did not probe the culture medium for the specific factors that may be influencing the phenotypic switch, investigation of proteins in the Wnt pathway, such as Dickkopf 1 homolog (DKK1) and frizzled-related protein (FRZB), and their ensuing effect on antagonizing the Wnt pathway, as others have shown [50], is an avenue of future research in examining the effects of viable bone on tissue engineered cartilage.

Taken together, the *in vivo* and *in vitro* studies presented here indicate that even a mild hypertrophic phenotype existing *in vitro* may allow for chondrogenically-primed tissue engineered constructs to undergo advanced hypertrophy and endochondral ossification. Conversely, *in vivo* implantation of acellular scaffolds or MSC-PCL constructs that were not precultured under chondrogenic conditions generated functional tissue with aggregate moduli of approximately 0.2 MPa without any detectable mineralization. This finding would suggest use of either “point of care” MSC seeding of the scaffolds (MSC-PCL) or simple implantation of acellular scaffolds (PCL implants) and reliance on endogenous cell infiltration to populate the scaffold to minimize this hypertrophic response. Importantly, substantial generation of repair tissue within the interstices and fiber bundles of the 3D woven scaffolds was demonstrated under either of the aforementioned conditions (MSC-PCL and PCL implants). While this repair tissue did not resemble hyaline-cartilage at the time of harvest, the harvested constructs possessed the functional compressive properties at the lower limits reported for native articular cartilage, even without significant GAG accumulation within the structure.

It is important to note that the use of the ectopic model to study the developing phenotype of an osteochondral construct may not be predictive of the same construct in an orthotopic model. From both our *in vivo* and *in vitro* data, it is clear that signals from soluble factors in the environment coupled with the initial construct phenotype affect the ultimate differentiation potential of a Ch-MSC-PCL tissue engineered construct. As a result, further study of chondrogenically primed MSCs in an orthotopic model is needed to elucidate the role of the local environment and mechanical forces on the ultimate phenotype of the repair tissue. Along these lines, a previous orthotopic study of stem cells in knees of mini-pigs showed that appropriate signaling molecules and mechanical stimuli are required for stem cell differentiation into non-hypertrophic chondrocytes [24]. Additionally, as previously alluded to, current MSC chondrogenesis protocols that have been successful at driving chondrogenesis *in vitro* may prove inadequate to sufficiently prepare the cells for *in vivo* use, and, therefore, improved differentiation protocols and/or improved tissue development *ex vivo* may be required to develop more stable tissue for implantation. Another area of active research involves devising more persistent chondrogenic, anti-hypertrophic, and/or anti-osteogenic cues *in vivo*, as will be required to repair osteoarthritic-associated lesions or other cartilage pathologies. As such, techniques such as combining gene therapy with tissue engineering may be used to deliver appropriate morphogens or cytokines protect the tissue once implanted from not only inflammation but also to provide factors that may push the cells to a desired osteogenic or chondrogenic phenotype [65-67].

## Acknowledgments

We thank EMS/Griltech for providing PCL yarn. We are grateful to C. Whittaker and M.E. Kolewe for many useful discussions and to X. Yin for critically reviewing the manuscript. This work was supported in part by the National Institutes of Health (NIH) (Grant Numbers R42 AR055404, to L.E.F and Cytex Therapeutics, Inc., P41 EB021911, P30 AR057235), the AO Foundation, and the Arthritis Foundation. The contents of this manuscript are solely the responsibility of the authors and do not necessarily represent the official views of the NIAMS or the NIH. Additional support for the Swanson Biotechnology Center Genomics and Bioinformatics and Computing Core Facilities at the MIT’s Koch Institute was from the National Cancer Institute (NCI) (Grant Number P30CA014051).

## Author contributions

B.L.L., L.E.F., and F.G. conceived and designed the project. B.L.L., S.N.Y., H.P., F.T.M., P.B.W., and J.F.W., performed the experiments. B.L.L., B.T.E., C.J.B, and L.E.F. analyzed the data and wrote the manuscript, with contributions from R.L., J.F.W., and F.G.

## Data and materials availability

Data and materials not presented in the article are available from the authors.

## Supplementary data

**Figure S1.**
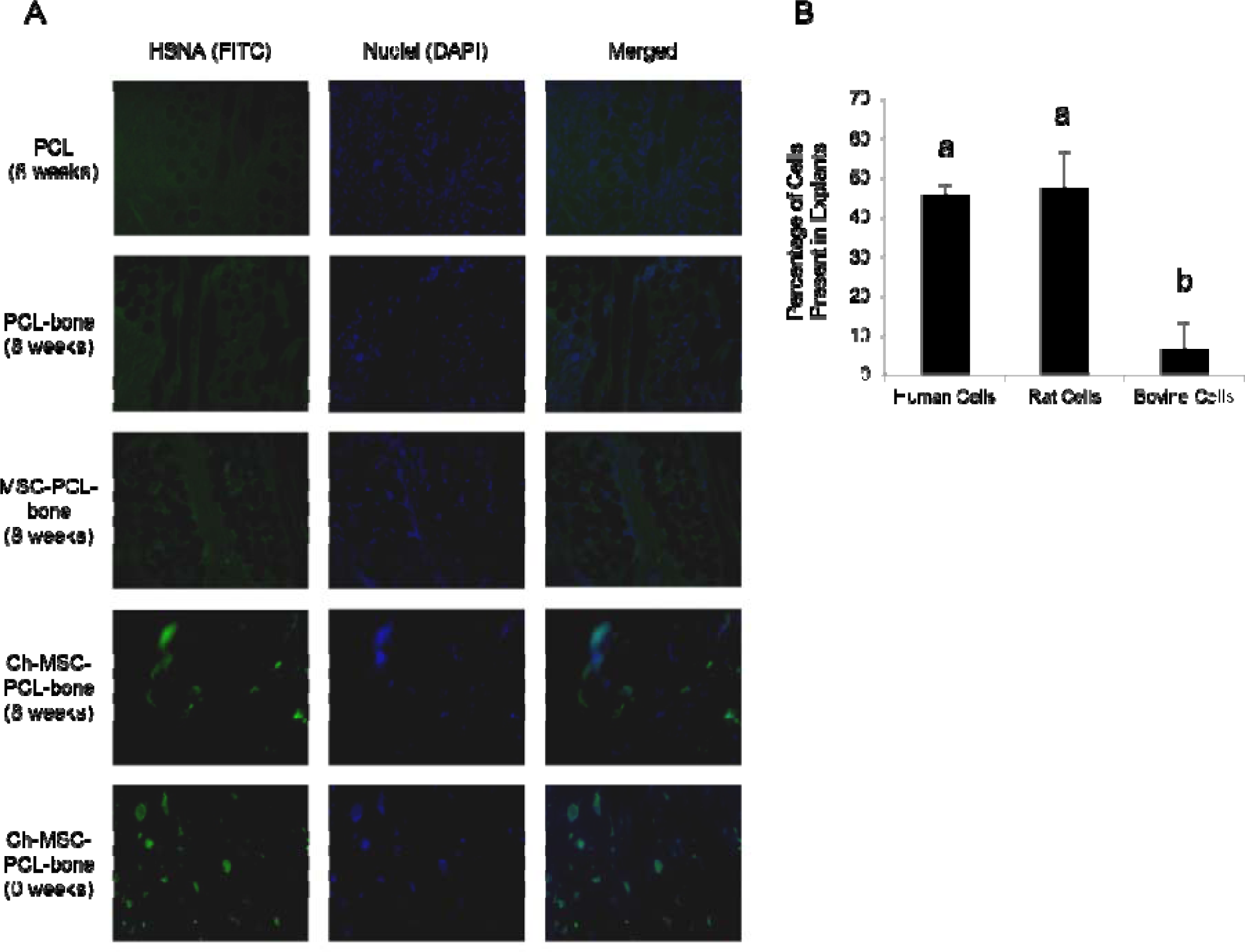
Cellular origin in implants after 8 weeks in vivo. A) Human cells, visualized by staining histological specimens of implants and explants for HSNA. The HSNA appears green (FITC); nuclei are counterstained blue (DAPI). (B) Relative contributions of each species to total RNA in the Ch-MSC-PCL bone group, assessed by RNA-seq transcriptome profile analyses. Groups not sharing the same letters are statistically different (p < 0.05).

